# Drivers of genetic diversity in secondary metabolic gene clusters within a fungal species

**DOI:** 10.1101/149856

**Authors:** Abigail L. Lind, Jennifer H. Wisecaver, Catarina Lameiras, Philipp Wiemann, Jonathan M. Palmer, Nancy P. Keller, Fernando Rodrigues, Gustavo H. Goldman, Antonis Rokas

## Abstract

Filamentous fungi produce a diverse array of secondary metabolites (SMs) critical for defense, virulence, and communication. The metabolic pathways that produce SMs are found in contiguous gene clusters in fungal genomes, an atypical arrangement for metabolic pathways in other eukaryotes. Comparative studies of filamentous fungal species have shown that SM gene clusters are often either highly divergent or uniquely present in one or a handful of species, hampering efforts to determine the genetic basis and evolutionary drivers of SM gene cluster divergence. Here we examined SM variation in 66 cosmopolitan strains of a single species, the opportunistic human pathogen *Aspergillus fumigatus*. Investigation of genome-wide within-species variation revealed five general types of variation in SM gene clusters: non-functional gene polymorphisms, gene gain and loss polymorphisms, whole cluster gain and loss polymorphisms, allelic polymorphisms where different alleles corresponded to distinct, non-homologous clusters, and location polymorphisms in which a cluster was found to differ in its genomic location across strains. These polymorphisms affect the function of representative *A. fumigatus* SM gene clusters, such as those involved in the production of gliotoxin, fumigaclavine, and helvolic acid, as well as the function of clusters with undefined products. In addition to enabling the identification of polymorphisms whose detection requires extensive genome-wide synteny conservation (e.g., mobile gene clusters and non-homologous cluster alleles), our approach also implicated multiple underlying genetic drivers, including point mutations, recombination, genomic deletion and insertion events, as well as horizontal gene transfer from distant fungi. Finally, most of the variants that we uncover within *A. fumigatus* have been previously hypothesized to contribute to SM gene cluster diversity across entire fungal classes and phyla. We suggest that the drivers of genetic diversity operating within a fungal species shown here are sufficient to explain SM cluster macroevolutionary patterns.

## Introduction

Filamentous fungi produce a diverse array of small molecules that function as toxins, antibiotics, and pigments [1]. Though by definition these so-called specialized or secondary metabolites (SMs) are not strictly necessary for growth and development, they are critical to the lifestyle of filamentous fungi [2]. For example, antibiotic SMs give their fungal producers a competitive edge in environments crowded with other microbes [3]. SMs can additionally mediate communication between and within species, as well as contribute to virulence on animal and plant hosts in pathogenic fungi [4,5].

A genomic hallmark of SMs in filamentous fungi is that the biosynthetic pathways that produce them are typically organized into contiguous gene clusters in the genome [6]. These gene clusters contain the chemical backbone synthesis genes whose enzymatic products produce a core metabolite, such as non-ribosomal peptide synthases (NRPS) and polyketide synthases (PKS), tailoring enzymes that chemically modify the metabolite, transporters involved in product export, and often transcription factors that control the expression of the clustered genes [6]. These gene clusters also occasionally contain resistance genes that confer self-protection against reactive or toxic metabolites [6]. Filamentous fungal genomes, particularly those in the phylum Ascomycota [6], typically contain dozens of SM gene clusters. However, most individual SM gene clusters appear to be either species-specific or narrowly taxonomically distributed in only a handful of species [7,6]. SM gene clusters that are more broadly distributed show discontinuous taxonomic distributions and are often highly divergent between species. Consequently, the identity and total number of SM gene clusters can vary widely even between very closely related species whose genomes exhibit very high sequence and synteny conservation [8,9].

In the last decade, several comparative studies have described macroevolutionary patterns of SM gene cluster diversity. For example, studies centered on genomic comparisons of closely related species have identified several different types of inter-species divergence, from single nucleotide substitutions (e.g., differences in fumonisins produced by *Fusarium* species are caused by variants in one gene [10]), to gene gain / loss events (e.g., the trichothecene gene clusters in *Fusarium* species and the aflatoxin family SM gene clusters in *Aspergillus* species) [11–16], and genomic rearrangements (e.g., the trichothecene gene clusters in *Fusarium*) [11]. Additionally, genetic and genomic comparisons across fungal orders and classes have identified several instances of gene gain or loss [17–19] and horizontal gene transfer [13,20–23] acting on individual genes or on entire gene clusters, providing explanations for the diversity and discontinuity of the taxonomic distribution of certain SM gene clusters across fungal species.

Although inter-species comparative studies have substantially contributed to our understanding of SM diversity, the high levels of evolutionary divergence of SM clusters make inference of the genetic drivers of SM gene cluster evolution challenging; put simply, it has been difficult to “catch” the mechanisms that generate SM gene cluster variation “in the act”. Several previous studies have examined intra-species or population-level differences in individual SM gene clusters, typically focusing on the presence and frequency of non-functional alleles of clusters involved in production of mycotoxins. Examples of clusters exhibiting such polymorphisms include the gibberellin gene cluster in *Fusarium oxysporum* [24], the fumonisin gene cluster in *Fusarium fujikuroi* [25], the aflatoxin and cyclopiazonic acid gene clusters in *Aspergillus flavus* [26], and the bikaverin gene cluster in *Botrytis cinerea* [27]. While these studies have greatly advanced our understanding of SM gene cluster genetic variation and highlighted the importance of within-species analyses, studies examining the entirety of SM gene cluster polymorphisms within fungal species are so far lacking. We currently do not know the types and frequency of SM gene cluster polymorphisms within fungal species, whether these polymorphisms affect all types of SM gene clusters, or the genetic drivers of SM gene cluster evolution.

To address these questions, we investigated the genetic diversity of all 36 known and predicted SM gene clusters in whole genome sequence data from 66 strains, 8 of which were sequenced in this study, of the opportunistic human pathogen *Aspergillus fumigatus*, a species with cosmopolitan distribution and panmictic population structure [28]. We found that 13 SM gene clusters were generally conserved and harbored low amounts of variation. In contrast, the remaining 23 SM gene clusters were highly variable and contained one or more of five different types of genetic variation: single-nucleotide polymorphisms including nonsense and frameshift variants, individual gene gain and loss polymorphisms, entire cluster gain and loss polymorphisms, polymorphisms associated with changes in cluster genomic location, and clusters with non-homologous alleles resembling the idiomorphs of fungal mating loci. Many clusters contained interesting combinations of these types of polymorphisms, such as pseudogenization in some strains and entire cluster loss in others. The types of variants we find are likely generated by a combination of DNA replication and repair errors, recombination, genomic insertions and deletions, and horizontal transfer. We additionally find an enrichment for transposable elements (TEs) around horizontally transferred clusters, clusters that change in genomic locations, and idiomorphic clusters. Taken together, our results provide a guide to both the types of polymorphisms and the genetic drivers of SM gene cluster diversification in filamentous fungi. As most of the genetic variants that we observe have been previously associated with SM gene cluster diversity across much larger evolutionary distances and timescales, we argue that processes influencing SM gene cluster diversity within species are sufficient to explain SM cluster macroevolutionary patterns.

## Results and Discussion

We analyzed the genomes of 66 globally distributed strains of *Aspergillus fumigatus* for polymorphisms in SM gene clusters. We performed whole-genome sequencing on 8 strains, and collected the remaining 58 strains from publicly available databases including NCBI Genome and NCBI Short Read Archive (Figure 1, (Table S1) [28–32]. All publicly available strains of *A. fumigatus* with sequencing data passing quality thresholds (see Methods) or with assembled genomes were included in our analysis. The resulting dataset contains strains sampled from 12 sites world-wide and from clinical and environmental sources (Table S1).

**Figure 1.**
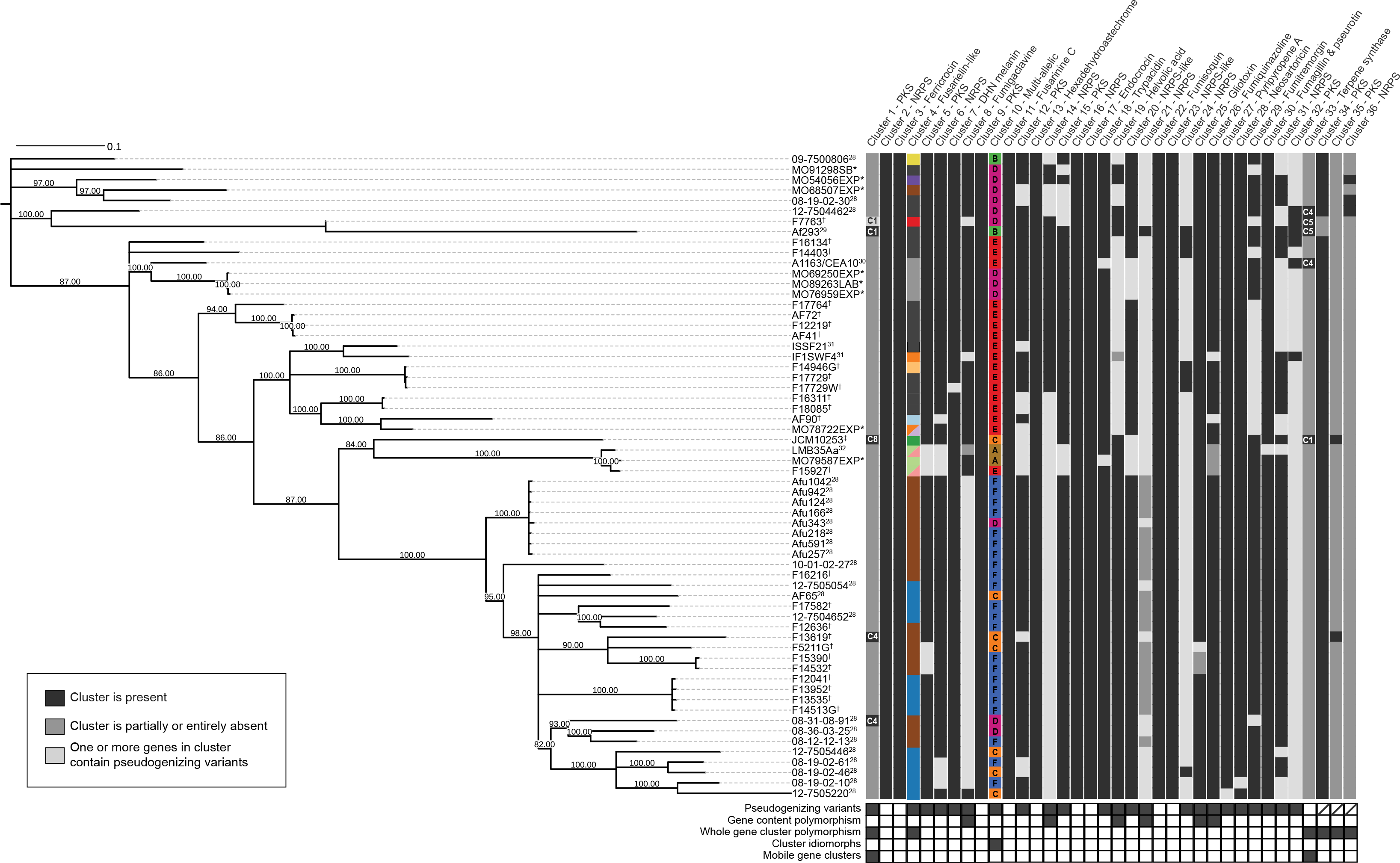
Genetic diversity of secondary metabolic gene clusters within a fungal species. The phylogeny was constructed using 15,274 biallelic SNPs with no missing data. The tree is midpoint rooted and all branches with bootstrap support less than 80% are collapsed. This phylogeny does not include strains Af10, Af210, Z5, or RP-2014 as short read data were not available. Superfixes following strain names indicate publications associated with DNA sequencing. * indicates strains sequenced in this study, † indicates strains sequenced at JCVI with no associated publication, and ‡ indicates strains sequenced by RIKEN with no associated publications. Heatmaps show presence, absence and polymorphisms in SM gene clusters. Black indicates the cluster is present in a strain with no polymorphisms aside from missense variants, light gray indicates one or more genes in the cluster are pseudogenized, and dark gray indicates the cluster is partially or entirely absent. Colors for Cluster 4 indicate which pseudogenizing variants are present (see Figure 3) and colors for Cluster 10 indicate which allele of the cluster is present (see Figure 4). Chromosomal location of Clusters 1 and 33 are indicated. If more than one type of polymorphism is present within a cluster in a strain, only one is depicted. Types of polymorphisms found in each cluster are summarized below the cluster heatmap.

We analyzed all strains for polymorphisms in 33 curated SM gene clusters present in the reference Af293 genome, and additionally searched for novel SM gene clusters (see Methods). These examinations revealed five distinct types of polymorphisms in SM gene clusters (Figure 1, Table 1):

**Table 1.** Types and rates of SM gene cluster variants in *A. fumigatus* strains.

| Description | Phenotype | Drivers | Frequency at cluster level | Frequency at strain level | Previous reports |
| --- | --- | --- | --- | --- | --- |
| Single-nucleotide polymorphisms and indels | Potential for protein function change (missense); abrogation of protein function (nonsense and frameshift) | DNA replication errors; relaxation of purifying selection | 100% (33/33 clusters; missense); 70% (23/33 clusters; nonsense and frameshift) | Every strain affected | Bikaverin in *Botrytis* [17,27], aflatoxin in *Aspergillus oryzae* and *Aspergillus flavus* [26], fumonisins in *Fusarium* [10], many others |
| Gene content polymorphisms | Loss of gene cluster function; structural changes in the metabolite; change in cluster expression or metabolite transport | Deletion and insertion events; recombination; transposable elements | 6 clusters | 27 / 66 strains | Trichothecene in *Fusarium*, aflatoxin and sterigmatocystin in *Aspergillus* [11–15], HC toxin in *Cochliobolus carbonarum* [96] |
| Whole gene cluster polymorphisms | Loss or gain of novel metabolites | Deletion and insertion events; horizontal gene transfer; transposable elements | 6 clusters | 13 / 66 strains | Gibberellin and fumonisin in *Fusarium* [24,25] |
| Cluster idiomorphs | Changes in metabolites produced or structure of metabolites | Transposable elements; recombination; other mechanisms? | 1 gene cluster | 8 unique identified alleles | Putative SM gene clusters in dermatophytes; putative SM gene cluster in *Aspergillus flavus* and *Aspergillus oryzae* [56,57] |
| Mobile gene clusters | Potential for change in gene regulation | Transposable elements; horizontal gene transfer; other mechanisms? | 2 gene clusters | 8 / 66 strains | None |

a) Single nucleotide and short indel polymorphisms. 33 / 33 SM gene clusters (present in the reference Af293 strain) contained multiple genes with missense SNPs and short indel variants in one or more strains. 23 / 33 SM gene clusters contained one or more genes with frameshift or nonsense variants.
b) Gene content polymorphisms involving loss or gain of one or more genes. 6 / 33 SM gene clusters contained a gene content polymorphism.
c) Whole SM gene cluster gain and loss polymorphisms. 3 / 33 SM gene clusters were entirely absent in one or more strains and an additional 3 previously unknown SM gene clusters were discovered.
d) Idiomorphic polymorphisms. One locus contained multiple non-homologous SM gene cluster alleles in different strains.
e) Genomic location polymorphisms. 2 / 33 SM gene clusters were found on different chromosomes between strains.

Both genomic location polymorphisms and idiomorphic polymorphisms are novel types of variants that have not been previously described for secondary metabolic gene clusters, likely because they can only be identified when genome-wide synteny and sequence conservation are high. The remaining types of variants, including single-nucleotide changes and gene gain and loss events, have been implicated at the species level as major drivers of secondary metabolic gene cluster evolution (Table 1), suggesting that the diversity-generating processes observed within a species are sufficient to explain SM gene cluster evolution across species.

### Single-nucleotide and indel polymorphisms

It is well established that single nucleotide polymorphisms (SNPs) and short indel polymorphisms, which are caused by errors in DNA replication and repair, are a major source of genomic variation [33]. Non-synonymous SNPs and indels with missense, frameshift, and nonsense effects were widespread across the 33 SM reference gene clusters (Figure 1, Table S2). Every strain contained numerous missense mutations and at least one nonsense or frameshift mutation in its SM gene clusters. Although missense mutations are likely to influence SM production, the functional effects of nonsense and frameshift mutations are comparatively easier to infer from genomic sequence data because they often result in truncated proteins and loss of protein function.

SNPs and short indel polymorphisms can affect secondary metabolite production, as in the case of the lack of trypacidin production in the A1163 strain because of a previously identified frameshift mutation in the polyketide synthase (PKS) of the trypacidin gene cluster [34]. Interestingly, we identified a premature stop codon (Gln273*) in a transcription factor required for trypacidin production, *tpcD* (Afu4g14550), in a strain sequenced in this study (M079587EXP) (Table S2). These data suggest that function of this SM gene cluster has been lost at least twice independently in *A. fumigatus*.

Individual nonsense or frameshift variants varied in frequency. For example, the non-ribosomal peptide synthase (NRPS) *pes3* gene (Afu5g12730) in SM gene cluster 21 harbors 16 nonsense or frameshift polymorphisms in 55 strains, seven of which are common (present in ≥10 strains) and another seven are rare (≤5 strains). Strains with lab-mutated null alleles of the *pes3* gene are more virulent than strains with functional copies [35], which may explain the widespread occurrence of null *pes3* alleles within *A. fumigatus*.

### Gene content polymorphisms

We additionally identified several SM gene clusters that gained or lost genes in some strains. These gene content polymorphisms were most likely generated through genomic deletion or insertion events and were sometimes found at high frequencies among strains (Figure 1, Table 1). In three cases, these polymorphisms impact backbone synthesis genes, rendering the SM gene cluster non-functional.

One example involves SM gene cluster 14, whose standard composition includes a pyoverdine synthase gene, an NRPS-like gene, an NRPS backbone gene, and several additional modification genes (Figure 2A). Four of the 66 strains examined lack an 11-kb region on the 3’ end of the cluster, which normally contains an NRPS gene and two additional cluster genes, and the first non-SM genes on the 3’ end flanking the cluster. All *A. fumigatus* strains contain a *copia* family TE [36,37] at the 3’ end of the cluster, suggesting that TEs may have been involved in the generation of this polymorphism. While this polymorphism could have arisen through a deletion event, a homologous cluster lacking the 11-kb region is also present in the reference genomes of *Aspergillus lentulus* and *Aspergillus fischeri*, close relatives of *A. fumigatus* (Figure 2A). The most parsimonious explanation is that the genome of the *A. fumigatus* ancestor contained an SM gene cluster that lacked the 11-kb region, and that this genomic region was subsequently gained and increased in frequency within *A. fumigatus*.

**Figure 2.**
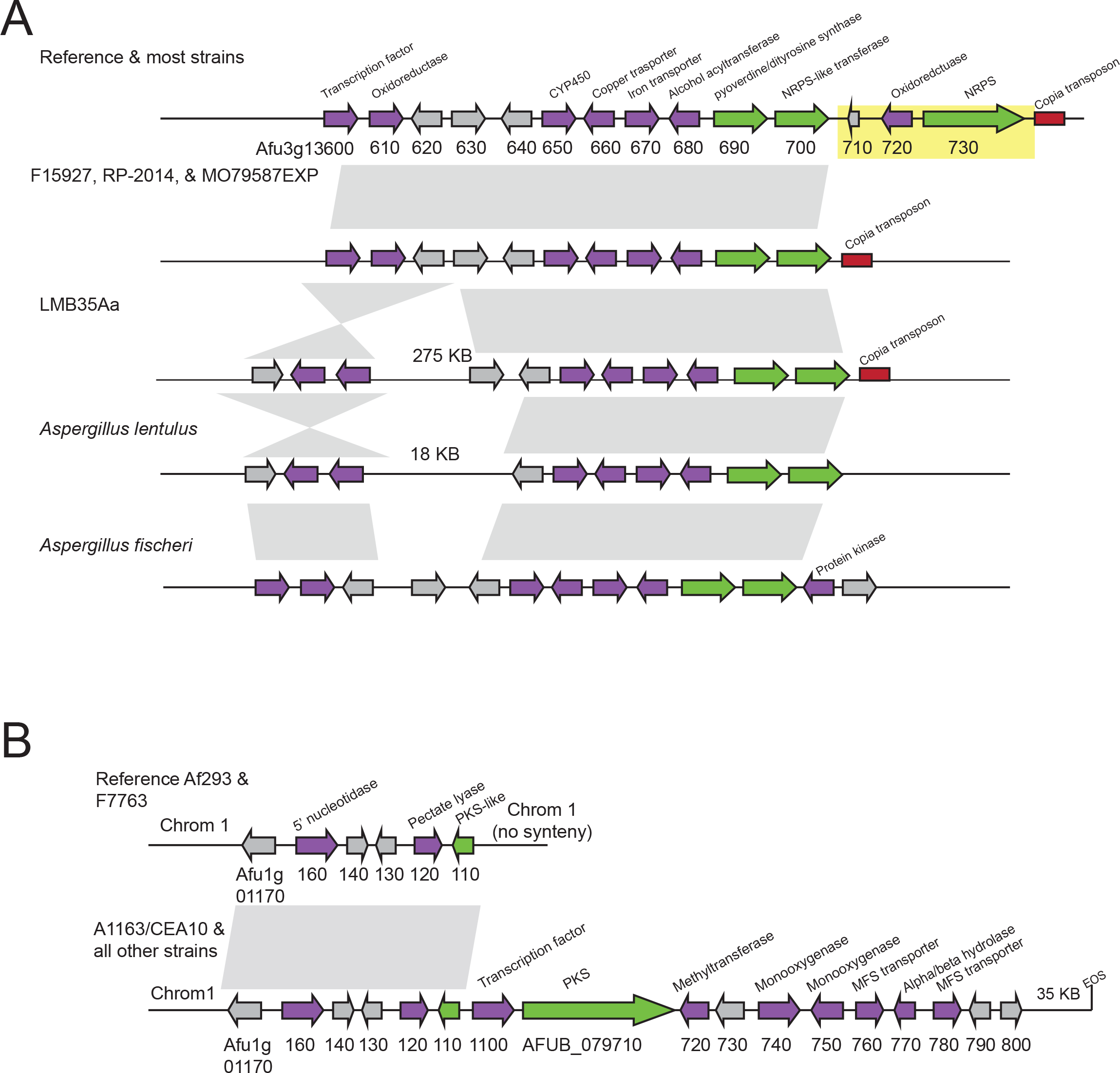
Gene gains and deletions in SM gene clusters. (A) Differences in gene content in SM gene cluster 14 in A. fumigatus strains and closely related species. Four A. fumigatus strains lack an 11-Kb region in this cluster, including an NRPS backbone gene, highlighted in yellow. Regions upstream and downstream of this cluster are syntenic. LMB35Aa also contains a large inversion that moves a transcription factor, oxidoreductase, and hypothetical protein 275 kb away from the cluster. Aspergillus fischeri and Aspergillus lentulus, close relatives of A. fumigatus, contain a cluster lacking the 11-kb region. (B) SM gene cluster found in most A. fumigatus strains but absent from the Af293 reference and from the F7763 strain. EOS denotes end of scaffold.

The remaining two gene content polymorphisms affecting SM backbone genes were restricted to one strain each and appear to have arisen through genomic deletion events. Specifically, strain IF1SWF4 lacks an 8-Kb region near the helvolic acid SM gene cluster, resulting in the loss of the backbone oxidosqualene cyclase gene as well an upstream region containing two non-SM genes (Figure S1). Strain LMB35Aa lacks a 54-kb region on the end of chromosome 2, which includes five genes from the telomere-proximal fumigaclavine C cluster (Figure S1).

Three other cases of gene content polymorphisms involved gene loss or truncation events of non-backbone structural genes. The second half of the ORF of the *gliM O*-methyltransferase gene in the gliotoxin gene cluster has been lost in 2 / 66 strains (Figure S1) and the first half of the permease *fmqE* in the fumiquinazoline gene cluster has been lost in 4 / 66 strains (Figure S1). Finally, an ABC transporter gene in SM cluster 21 has been almost entirely lost in 21 / 66 strains (Figure S1). This deletion event is found in strains that are related in the SNP-based strain phylogeny but does not perfectly mirror the phylogeny (Figure 1).

### Whole gene cluster loss polymorphisms

Several SM gene clusters were gained or lost entirely across strains. We observed several instances where a cluster present in the genome of either the reference Af293 or A1163 (also known as CEA10) strains was absent or pseudogenized in other strains, which we present in this section.

One of the novel SM gene clusters, cluster 34, was present in all but two of the strains (Af293 and F7763). Cluster 34 contains a PKS backbone gene, one PKS-like gene with a single PKS associated domain, nine genes with putative biosynthetic functions involved in secondary metabolism, and six hypothetical proteins (Figure 2B). The two strains that lack cluster 34 contain a likely non-functional cluster fragment that includes the PKS-like gene, two biosynthetic genes, and three hypothetical proteins. Interestingly, the 3’ region flanking cluster 34 is syntenic across all 66 strains but the 5’ region is not, suggesting that a recombination or deletion event may have resulted in its loss in the Af293 and F7763 strains. These two strains form a clade in the strain phylogeny (Figure 1), so it is likely that this deletion or recombination event occurred once.

One notable example of an SM gene cluster present in the Af293 reference genome but absent or pseudogenized in others was SM cluster 4. This cluster contains 5 genes on the tip of the Af293 chromosome 1, and contains orthologs to five of the six genes in the fusarielin-producing gene cluster in *Fusarium graminearum* [38]. Cluster 4 is also present in several other *Aspergillus* species, including *A. clavatus* and *A. niger* [38], as well as in whole or in part in other *non-Aspergillus* fungi in the class Eurotiomycetes and in fungi in the class Sordariomycetes (Figure S3) [30,39–47]. Phylogenetic analysis of the genes in cluster 4 does not provide a clear view of the origin of this cluster, which is consistent either with extensive gene loss in both Sordariomycetes and Eurotiomycetes, or alternatively with horizontal gene transfer (HGT) between fungi belonging to the two classes (Figure S2, Figure S3).

Cluster 4 is entirely absent in 4 / 66 strains, and its genes are undergoing pseudogenization in an additional 43 strains via multiple independent mutational events (Figure 3). The four strains lacking the cluster form a single clade on the strain phylogeny, suggesting that the cluster was lost in a single deletion event (Figure 1). Further, 19 strains shared a single frameshift variant in the polyketide synthase gene (4380_4381insAATGGGCT; frameshift at Glul461 in Afulgl7740) and an additional 13 strains shared a single frameshift variant (242delG; frameshift at Gly81) in an aldose 1-epimerase gene (Afulgl7723) (Figure 3A, Table S2). Eleven other strains each contained one to several frameshift or nonsense polymorphisms involving nine unique mutational sites. Five of these strains contained multiple distinct frameshifts and premature stop codons in more than one gene in the cluster, indicating that the entire pathway is pseudogenized in these strains.

**Figure 3.**
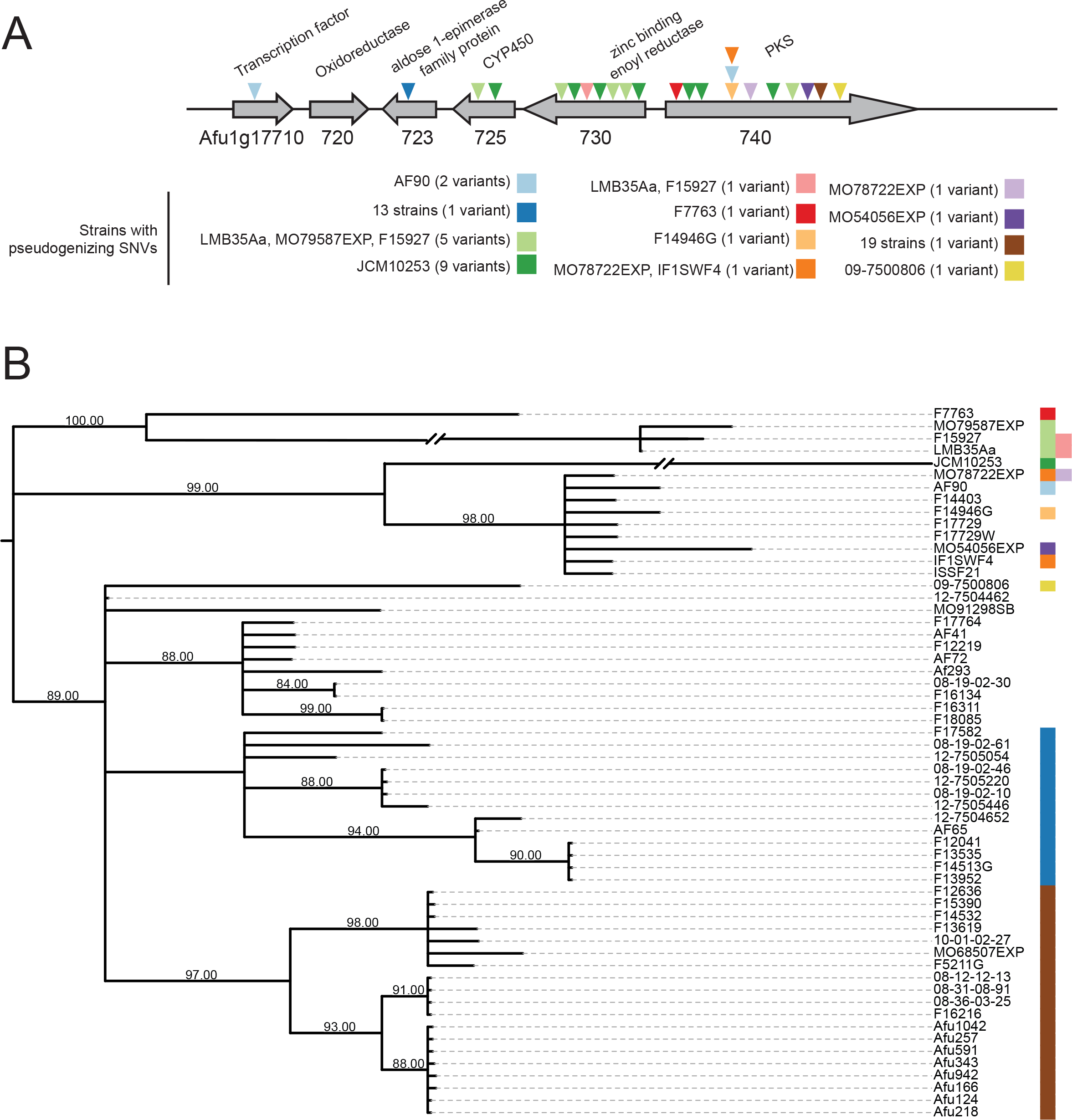
Pseudogenization in the fusarielin-like SM gene cluster. (A) Positions of frameshift variants and nonsense variants in the fusarielin-like SM gene cluster 4. (B) Locus phylogeny of the fusarielin-like SM gene cluster based on a nucleotide alignment of the entire gene cluster including intergenic and non-coding regions. The phylogeny is midpoint rooted and branches with bootstrap support <80% are collapsed. Two branches were shortened for visualization purposes. Strains with pseudogenizing variants are indicated with colored boxes. Colors correspond to variants shown in (A).

A phylogeny of the entire cluster 4 locus across all 62 strains with short-read data shows that two pseudogenizing variants shared across multiple strains, one in the aldose 1-epimerase gene and one in the polyketide synthase, are found in loci that form well-supported clades (Figure 3B), suggesting that these variants arose once. Similarly, a set of variants shared across three strains and one variant shared in two strains are found in loci that form well-supported clades in the locus phylogeny. Two strains sharing a pseudogenizing variant in the polyketide synthase do not group together in the locus phylogeny, a discordance likely stemming from within-locus recombination events. Finally, functional alleles of cluster 4 are distributed throughout the locus phylogeny, suggesting that the functional allele is ancestral and the pseudogenized variants are derived.

Perhaps surprisingly, loss of function polymorphisms (from nonsense and frameshift mutations to wholesale cluster loss) are common and sometimes frequent within *A. fumigatus*. The majority of these polymorphisms are presumably neutral, and reflect the fact that any mutation is more likely to result in loss of a function than in gain. Consistent with this hypothesis is our observation that these loss events were often found at low frequencies. However, the possibility also exists that some of the high-frequency, recurrent loss of function polymorphisms may be adaptive. Given that many secondary metabolites are primarily secreted in the extracellular environment and can benefit nearby conspecifics that are not themselves producing the metabolite [51], individual strains may be circumventing the energetically costly process of producing the metabolite themselves in a situation analogous to the Black Queen hypothesis [52].

### Whole gene cluster gain polymorphisms

By searching for novel SM gene clusters in the genomes of the other 65 *A. fumigatus* strains, we found three SM gene clusters that were absent from the genome of the Af293 reference strain. As SM gene clusters are often present in repeat-rich and subtelomeric regions that are challenging to assemble [48,49], the strains analyzed here might harbor additional novel SM gene clusters that were not captured here. One of these SM gene clusters, cluster 34, was mentioned earlier as an example of whole gene cluster loss polymorphism (Figure 2B) and is present in most strains but has been lost in two strains. The other two SM gene clusters absent from the Af293 genome are present at lower frequencies and likely reflect gene cluster gain events; cluster 35 is present in 2 / 66 strains and cluster 36 in 4 / 66 strains. Cluster 35 is located in a region syntenic with an Af293 chromosome 4 region and is flanked on both sides by TEs (Figure S4). Eight of the 14 genes in this SM gene cluster are homologous to genes in an SM gene cluster in the genome of the insect pathogenic fungus *Metarhizium anisopliae* (Figure S4) [50]. Phylogenetic analysis of these 8 genes is consistent with a horizontal transfer event (Figure S5). The two strains that contain this novel cluster are not sister to each other on the strain phylogeny (Figure 1).

Cluster 36 is an NRPS-containing cluster located on shorter genomic scaffolds that lack homology to either the Af293 or A1163 genomes, making it impossible to determine on which chromosome this cluster is located (Figure S4). Two of the strains containing this novel cluster are sister to each other on the strain phylogeny, while the third is distantly related to these two (Figure 1). The evolutionary histories of the genes in the cluster are consistent with vertical inheritance, and these genes are present in multiple *Aspergillus* species.

### Idiomorph polymorphisms

One of the most peculiar types of polymorphisms that we identified is a locus containing different unrelated alleles of SM gene clusters, reminiscent of the idiomorph alleles at the fungal mating loci [53]. This locus, which resides on chromosome 3 and corresponds to cluster 10 in the Af293 genome (Figure 4), was previously described as being strain-specific in a comparison between Af293 and A1163 strains [30] and is thought to reside in a recombination hot spot [28]. Our analysis showed that there are at least 6 different alleles of this cluster in *A. fumigatus*, containing 4 different types of key enzymes involved in natural product biosynthesis: a polyketide synthase (PKS) non-ribosomal peptide synthetase (NRPS) hybrid, a highly reducing (HR) PKS, a non-reducing (NR) PKS and an NRPS-like enzyme (Figure 4). Two additional alleles were present in only one strain each (Figure S6).

**Figure 4.**
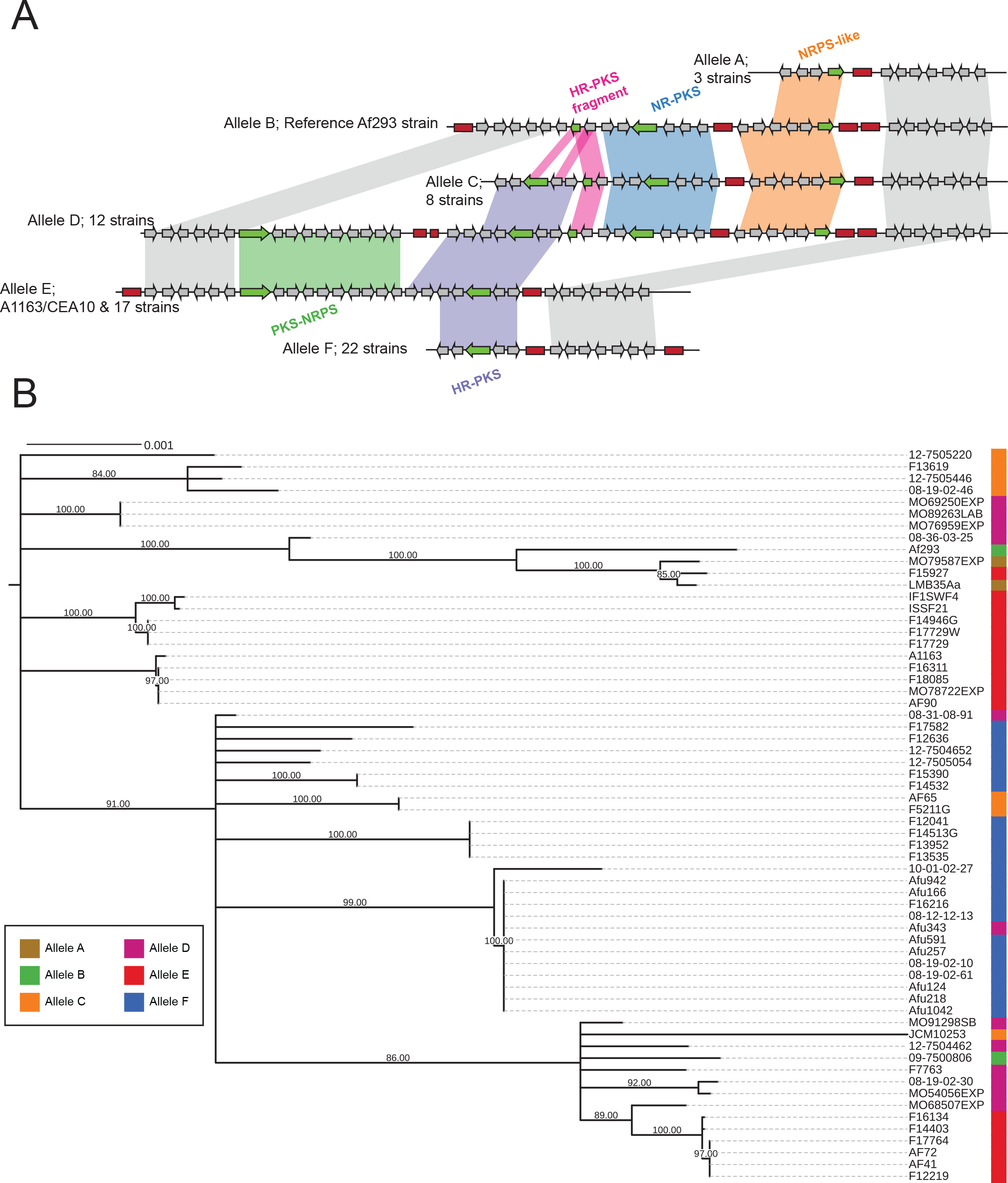
Six alleles of an idiomorphic SM gene cluster. (A) Alleles of SM gene cluster 10 on chromosome 3. Red boxes denote transposable elements. Green arrows denote backbone genes (PKS or NRPS). (B) Locus phylogeny of conserved downstream of the idiomorph cluster (highlighted in gray in A). Phylogeny was constructed using a 48 MB nucleotide alignment with the GTRGAMMA model and midpoint rooted. Branches with bootstrap support < 80% were collapsed.

In the Af293 reference genome, the cluster present at the idiomorph locus contains one NR-PKS along with an NRPS-like gene (Allele B). In the A1163 reference genome and 17 other strains, there is a PKS-NRPS and an HR-NRPS at this locus (Allele E). These alleles show an almost complete lack of sequence similarity except for a conserved hypothetical protein and a fragment of the HR-PKS in the Af293 allele; in contrast, the upstream and downstream flanking regions of the two alleles, which do not contain any backbone genes, are syntenic. Remarkably, another allele, present in 12 strains, contains all of the genes from both the Af293 and A1163 clusters (Allele D). The remaining three alleles contain various combinations of these genes. One allele found in 22 strains contains some A1163-specific genes, including the HR-PKS, and no Af293-specific genes (Allele F), while another allele found in 3 strains contains some Af293-specific genes, including the NRPS-like gene, but no A1163 genes (Allele A). The final allele, present in 8 strains, contains the entire Af293 allele as well as part of the A1163 allele containing the HR-PKS (Allele C). Every allele is littered with multiple long terminal repeat sequence fragments from *gypsy* and *copia* TE families as well as with sequence fragments from DNA transposons from the *mariner* family [36]. In some cases, these TEs correspond with breakpoints in synteny between alleles, suggesting that the diverse alleles of this SM gene cluster may have arisen via TE-driven recombination. Further, both of the alleles that are restricted to a single strain have an insertion event of several genes near a TE, while the rest of the locus is highly similar to one of the more common alleles (Figure S6).

Untargeted XCMS analysis [54] of an Allele D strain (08-19-02-30) and two Allele F strains (08-12-12-13 and 08-19-02-10) and comparison of their metabolite profiles revealed the presence of 2 unique masses in 08-19-02-30 (Table S4; Figure S7), raising the possibility that variation at the idiomorph locus is functional. Further analysis is underway to investigate whether any of these m/z can be directly linked to the Allele D sequence.

To gain insight into the evolutionary history of this locus, we constructed a phylogeny based on its conserved downstream flanking region (Figure 4B). The resulting phylogeny shows some grouping of strains that share alleles, but there are no clades that contain all instances of a particular allele. This is likely to be the consequence of within-locus recombination between strains of *A. fumigatus*, which has been previously described at this locus [28] and which is potentially driven by the high number of repetitive sequences at this locus.

While it is tempting to speculate that Allele D, the longest allele containing all observed genes, represents the ancestral state, this does not explain the presence of a shared hypothetical protein and PKS gene fragment between Allele C and Allele B. Further, two close relatives of *A. fumigatus, A. lentulus* and *A. fischeri*, contain a similar region with conserved upstream and downstream flanking genes that is highly dissimilar from any of the alleles observed in *A. fumigatus* (Figure S8). In both species, this locus contains numerous TEs as well as genes homologous to portions of allele E in *A. fumigatus* (Figure S8). *A. fischeri* additionally contains two hypothetical proteins from the PKS-NRPS region of *A. fumigatus* and an additional hybrid PKS-NRPS-containing gene cluster not found in either *A. lentulus* or *any A. fumigatus* strain (Figure S8). Other genes at this locus in both *A. lentulus* and *A. fischeri* have functions likely not related to SM. Interestingly, *A. lentulus* contains a gene with a heterokaryon incompatibility protein domain, which may be involved in determining vegetative incompatibility [55]. Only one representative genome from each species has been sequenced, but based on the high concentration of TEs and lack of sequence similarity with any *A. fumigatus* alleles, it is likely that this locus is highly variable within both *A. lentulus* and *A. fischeri*.

It is possible that polymorphism at this locus originated via SM gene cluster fusion or fission events driven by TEs, which are present in large numbers. Interestingly, two other previously described instances of SM gene cluster variation bear some resemblance to the *A. fumigatus* idiomorphic SM gene cluster 10 locus. The first is the presence of two non-homologous *Aspergillus flavus* alleles, where some strains contain a 9-gene sesquiterpene-like SM gene cluster and others contain a non-homologous 6-gene SM gene cluster at the same genomic location [56]. The second is the presence of two non-homologous SM gene clusters at the same, well-conserved, locus in a comparison of six species of dermatophyte fungi [57]. Based on these results, we hypothesize that idiomorphic clusters may be common in fungal populations and contribute to the broad diversity of SM gene clusters across filamentous fungi.

### Genomic location polymorphisms

The final type of polymorphism that we observed is associated with SM gene clusters that are found in different genomic locations in different strains, suggesting that these SM gene clusters are behaving like mobile genetic elements. This type of polymorphism was observed in SM gene clusters 1 and 33, both of which produce as yet identified products, and are present at low frequencies in *A. fumigatus* strains.

SM gene cluster 1, which is present in six strains at three different genomic locations (Figure 5A), consists of a PKS and four other structural genes that are always flanked by a 15 Kb region (upstream) and a 43 Kb region (downstream) containing TEs. In the reference Af293 strain and in strain F7763, cluster 1 and its flanking regions are located on chromosome 1, while in strains 08-31-08-91, F13619, and Z5 they are located between Afu4g07320 and Afu4g07340 on chromosome 4. In contrast, in strain JCM10253, the cluster and flanking regions are located on chromosome 8 immediately adjacent to the 3′ end of the intertwined fumagillin and pseurotin SM gene supercluster [58]. The strains containing the allele on Chromosome 1 are sister to each other on the strain phylogeny, while the other strains are scattered across the tree and do not reflect the phylogeny (Figure 1).

**Figure 5.**
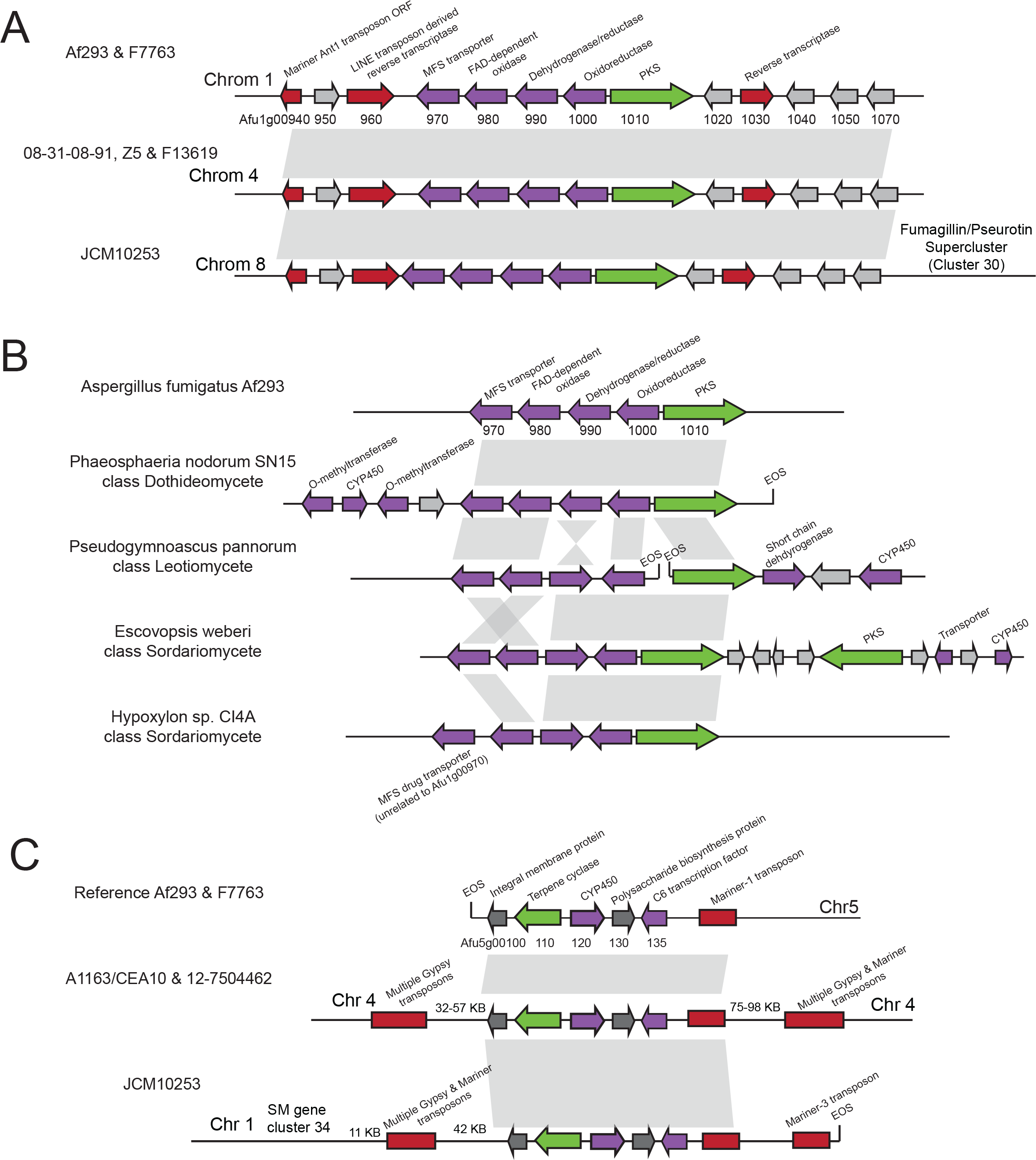
Multiple genomic locations of two SM gene clusters. (A) SM gene cluster 1 (Afu1g00970-01010) and flanking region is found in different genomic locations.The flanking regions contain transposon-derived open reading frames including two putative reverse transcriptases. In one strain, SM gene cluster 1 is found adjacent to SM gene cluster 30. (B) Synteny of A. fumigatus SM gene cluster 1 with clusters in *Phaeosphaeria nodorum, Pseudogymnoascus pannorum, Escovopsis weberi*, and *Hypoxylon sp. CI4A*.EOS denotes end of scaffold. All species contain non-syntenic genes predicted by antiSMASH to be part of a biosynthetic gene cluster. (C) SM gene cluster 33 (Afu5g00100-00135) is found in different genomic locations in diferent strains. In one strain, the cluster is adjacent to SM gene cluster 34. Multiple transposable elements flank the cluster in each strain.

In 5 / 6 strains, cluster 1 appears to be functional and does not contain nonsense SNPs or indels. However, the cluster found on chromosome 1 in strain F7763 contains two stop codons in the oxidoreductase gene (Glnl21* and Gln220*) and two premature stop codons in the polyketide synthase (Glnll56*) and Glnl542*)), suggesting this strain contains a null allele.

This “jumping” gene cluster is not present in any other sequenced genome in the genus *Aspergillus*, and phylogenetic analysis of its constituent genes is consistent with HGT between fungi (Figure S7). Specifically, this gene cluster is also present in *Phaeosphaeria nodorum* [59], a plant pathogen from the class Dothideomycetes, *Pseudogymnoascus pannorum* [60], a fungus isolated from permafrost from the Leotiomycetes, and *Escovopsis weberi* [61], a fungal parasite of fungus-growing ants from the Sordariomycetes (Figure 5B). One additional species, the endophyte *Hypoxylon* sp. CI4A from the class Sordariomycetes [62], contains four of the five cluster genes but is missing Afu1g00970, an MFS drug transporter. However, this species contains a gene unrelated to Afu1g00970 that is annotated as an MFS drug transporter immediately adjacent to this cluster (Figure 5B). None of these fungi contain the upstream or downstream TE-rich flanking regions present in *A. fumigatus*, and each fungus contains additional unique genes with putative biosynthetic functions adjacent to the transferred cluster. The most likely explanation for this change in flanking regions is that this SM gene cluster was transferred into *A. fumigatus* once and has subsequently “jumped” in different genomic locations in different strains.

The second SM gene cluster that shows variation in its genomic location across strains, cluster 33, contains a terpene synthase. This cluster is present in only 5 strains at 3 distinct locations (Figure 5C). Similar to cluster 1, cluster 33 is also flanked by TEs, and in one strain the cluster is located in a new region 58 Kb from SM gene cluster 34. Two strains that contain the cluster in the same genomic location are sister to each other on the strain phylogeny, while the placement of the other three strains containing the cluster does not reflect the phylogeny (Figure 1). In contrast to cluster 1, cluster 33 does not appear to have been horizontally transferred between fungi and its genes are present in other sequenced *Aspergillus* species [63], suggesting that the mobility of clusters 1 and 33 may be driven by different mechanisms.

Interestingly, both cases of mobile gene clusters are located near or immediately adjacent to other SM gene clusters in some strains. Cluster 33 is located 58Kb away from cluster 34 in one strain and Cluster 1 is located immediately adjacent to the intertwined fumagillin and pseurotin supercluster [58] in another. This supercluster is regulated by the transcriptional factor *fapR* (Afu8g00420) and is located in a chromosomal region controlled by the master SM regulators *laeA* (Afu1g14660) and *veA* (Afu1g12490) [58,64], raising the hypothesis that mobile gene clusters might be co-opting the regulatory machinery acting on adjacent SM gene clusters. Previous work has hypothesized that the fumagillin and pseurotin supercluster formed through genomic rearrangement events placing the once-independent gene clusters in close proximity to each other [58]. Our observation that the mobile cluster 1 is located in this same region not only supports this hypothesis but also implicates TEs as one of the mechanisms by which superclusters are formed. These superclusters may also represent an intermediate stage in the formation of new SM gene clusters. Supercluster formation, potentially mediated by mobile gene clusters, and followed by gene loss, could explain macroevolutionary patterns of SM gene clusters where clustered genes in one species are found to be dispersed over multiple gene clusters in other species [9,11].

### Conclusions

Our examination of the genomes of 66 strains of *Aspergillus fumigatus* revealed five general types of polymorphisms that describe variation in SM gene clusters. These polymorphisms include variation in SNPs and short indels, gene and gene cluster gains and losses, non-homologous (idiomorph) gene clusters at the same genomic position, and mobile clusters that differ in their genomic location across strains (Figure 6). Previous work has demonstrated that SM gene clusters, like the metabolites that they produce, are highly divergent between fungal species [8,9,19,63]. Our examination of genome-wide variation shows that these SM gene clusters are also diverse across strains of a single fungal species. These results also demonstrate that the diversity of SM gene clusters within *A. fumigatus* cannot be captured by sequencing a single representative strain, which is the current standard practice for determining the SM gene cluster content of a fungal species.

**Figure 6.**
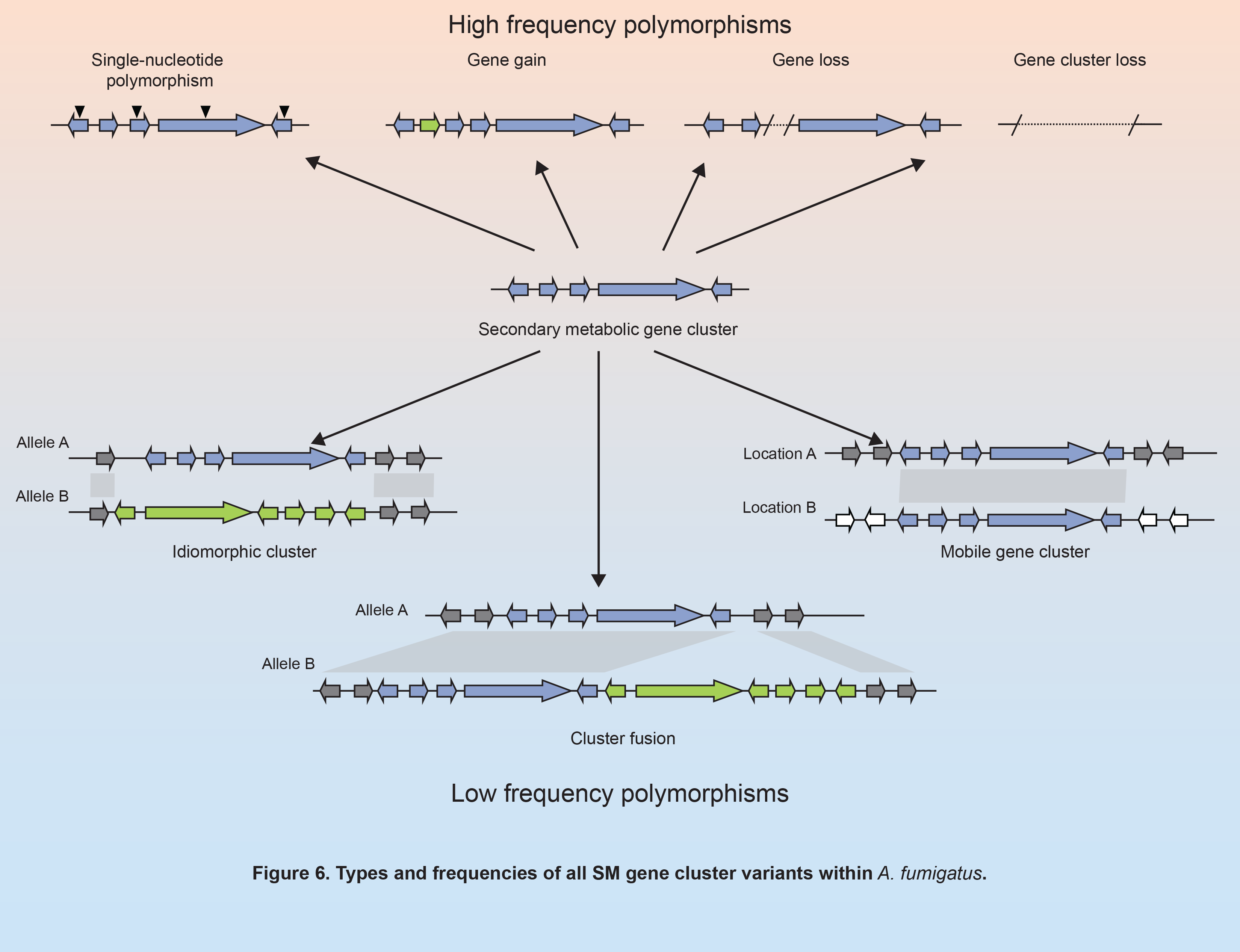
Types and frequencies of all SM gene cluster variants within. *A. fumigatus*.

The quantification of diversity in SM gene clusters within a species is dependent on both numbers and types of strains analyzed. The types of polymorphisms detected as well as their observed frequency, especially for rare polymorphisms, will increase with the number of genomes examined. In addition, both the frequencies of the different types of polymorphisms and the polymorphisms themselves may also change with sampling design, or in a manner corresponding to the population structure or ecology of the species under study. *A. fumigatus* is a cosmopolitan species with panmictic population structure [28], characteristics that do not always apply to other filamentous fungi. Fungi exhibiting strong population structure or fungi adapted to different ecological niches might contain different patterns of genetic diversity.

Nevertheless, the variants and genetic drivers we observe at the within-species level are also implicated as driving SM gene cluster variation at the between-species level, suggesting that the observed microevolutionary processes are sufficient to explain macroevolutionary patterns of SM gene cluster evolution. For example, the narrow and discontinuous distribution of SM gene clusters across the fungal phylogeny has been attributed to HGT as well as to gene cluster loss [13,15,20,22,30,65–67]. Here, we find evidence that both processes also influence the distribution of SM gene clusters within a species (Figures 2, 5, S2-S5). Interestingly, the fraction of SM gene clusters within *A. fumigatus* that harbor loss of function polymorphisms is substantial, consistent with the macroevolutionary view that SM gene cluster loss is rampant [18,19,67]. However, our within-species observations are also consistent with the macroevolutionary importance of HGT to SM gene cluster evolution. Once thought to be non existent in eukaryotes, HGT is now considered to be responsible for the presence of several different SM gene clusters in diverse filamentous fungi [13,67,68]. The instances of HGT of SM gene clusters within *A. fumigatus* suggests that acquisition of foreign genetic material containing SM gene clusters is likely a common and ongoing occurrence in fungal populations.

One recurring theme across different types of SM gene cluster polymorphisms in *A. fumigatus* was the perpetual presence of TEs adjacent to or within clusters. One particularly striking case is the “idiomorphic” Cluster 10, where TEs seem to correspond with breakpoints in synteny both within *A. fumigatus* and also between *A. fumigatus* and its close relatives (Figure 4, Figure S8). TEs were also present flanking mobile and horizontally transferred SM gene clusters and were located adjacent to gene gain sites. There are several potential explanations for the observed TE enrichment. First, TE presence may promote repeat-driven recombination and gene rearrangement, or the TEs themselves may be the agents of horizontally transferred clusters (either on their own or through a viral vector). Alternatively, it may simply be the case that SM gene clusters preferentially reside in TE-rich genomic regions.

In summary, examination of SM gene cluster variation within a single fungal species revealed five distinct types of polymorphism that are widespread across different types of SM gene clusters and are caused by many underlying genetic drivers, including errors in DNA transcription and repair, non-homologous recombination, gene duplication and loss, and HGT. The net effect of the observed variation raises the hypothesis that the chemical products of filamentous fungal species are in a state of evolutionary flux, each population constantly altering its SM gene cluster repertoire and consequently modifying its chemodiversity.

## Methods

### Strains analyzed

Eight strains of *A. fumigatus* were isolated from four patients with recurrent cases of aspergillosis in the Portuguese Oncology Institute in Porto, Portugal. Each strain was determined to be *A. fumigatus* using macroscopic features of the culture and microscopic morphology observed in the slide preparation from the colonies with lactophenol solution [69]. Based on the morphological characterization, all clinical strains were classified as *A. fumigatus complex-*Fumigati. After whole genome sequencing, retrieval and examination of the beta tubulin and calmodulin sequences of each strain confirmed that all strains belonged to *A. fumigatus* (see Phylogenetic Analysis and Figure S9). The genomes of all eight strains were sequenced using 150bp illumina paired-end sequence reads at the Genomic Services Lab of Hudson Alpha (Huntsville, Alabama, USA). Genomic libraries were constructed with the illumina TruSeq library kit and sequenced on an illumina HiSeq 2500 sequencer. Samples of all eight strains were sequenced at greater than 180X coverage or depth (Table S1). Short read sequences for these 8 strains are available in the Short Read Archive under accession SRP109032.

In addition to the 8 strains sequenced in this study, we retrieved 58 *A. fumigatus* strains with publicly available whole genome sequencing data, resulting in a dataset of 66 strains (Table S1). The strains used included both environmental and clinical strains and were isolated from multiple continents. Genome assemblies for 10 of these strains, including the Af293 and A1163 reference strains, were available for download from GenBank [28–32,70]. For 6 of these strains, short read sequences were also available from the NCBI Short Read Archive (SRA), which were used for variant discovery only (see Single nucleotide variant (SNV) and indel discovery) and not for genome assembly. Short read sequences were not available for the remaining 4 strains. Short read sequences were downloaded for an additional 48 strains from the Short Read Archive if they were sequenced with paired-end reads and at greater than 30× coverage.

### Single nucleotide variant (SNV) and indel discovery

All strains with available short read data (62 of 66 strains) were aligned to both the Af293 and A1163 reference genomes using BWA mem version 0.7.12-r1044 [71]. Coverage of genes present in the reference genome was calculated using bedtools v2.25.0 [72]. SNV and indel discovery and genotyping was performed relative to the Af293 reference genome and was conducted across all samples simultaneously using the Genome Analysis Toolkit version 3.5-0-g36282e4 with recommended hard filtering parameters [73–75] and annotated using snpEff version 4.2 [76].

### *De novo* genome assembly and gene annotation

All 56 strains without publicly available genome assemblies were *de novo* assembled using the iWGS pipeline [77]. Specifically, all strains were assembled using SPAdes v3.6.2 and MaSuRCA v3.1.3 and resulting assemblies were evaluated using QUAST v3.2 [78–80]. The average N50 of assemblies constructed with this strategy was 463 KB (Table S1). Genes were annotated in these assemblies as well as in five GenBank assemblies with no predicted genes using augustus v3.2.2 trained on *A. fumigatus* gene models [81]. Repetitive elements were annotated in all assemblies using RepeatMasker version open-4.0.6 [82].

### Secondary metabolic gene cluster annotation and discovery

Secondary metabolic gene clusters in the Af293 reference genome were taken from two recent reviews, both of which considered computational and experimental data to delineate cluster boundaries [83,84] (Table S3). The genomes of the other 65 strains were scanned for novel SM gene clusters using antiSMASH v3.0.5.1 [85]. To prevent potential assembly errors from confounding the analysis, any inference about changes in genomic locations of genes or gene clusters was additionally verified by manually inspecting alignments and ensuring that paired end reads supported an alternative genomic location (see SNV and indel discovery). Cases where paired end reads did not support the change in genomic location (i.e., all 3’ read mapping to Chromosome 1 and all 5’ pairs mapping to Chromosome 8), or where mapping was ambiguous or low quality were discarded.

### Phylogenetic analysis

To confirm all strains in this analysis belonged to the species *Aspergillus fumigatus*, the genomic sequences of the beta tubulin and calmodulin genes were extracted from the assembled genomes of all strains. Gene phylogenies were constructed using *Aspergillus fischerianus* as an outgroup using RA×ML v8.0.25 with the GTRGAMMA substitution model [86]. The tree was midpoint rooted and all branches with bootstrap support less than 80% were collapsed (Figure S10).

To construct a SNP-based strain phylogeny, biallelic SNPs with no missing data were pruned using SNPRelate v1.8.0 with a linkage disequilibrium threshold of 0.8 [87]. A total of 15,274 SNVs were used to create a phylogeny using RA×ML v8.0.25 with the ASC_BINGAMMA substitution model [86]. The tree was midpoint rooted and all branches with bootstrap support less than 80% were collapsed. The phylogeny was visualized using ITOL version 3.0 [88].

To understand the evolutionary histories of specific SM gene clusters showing unusual taxonomic distributions, we reconstructed the phylogenetic trees of their SM genes. Specifically, SM cluster protein sequences were queried against a local copy of the NCBI non-redundant protein database (downloaded May 30, 2017) using phmmer, a member of the HMMER3 software suite [89] using acceleration parameters --F1 1e-5 --F2 1e-7 --F3 1e-10. A custom peri script sorted the phmmer results based on the normalized bitscore (nbs), where nbs was calculated as the bitscore of the single best-scoring domain in the hit sequence divided by the best bitscore possible for the query sequence (i.e., the bitscore of the query aligned to itself). No more than five hits were retained for each unique NCBI Taxonomy ID. Full-length proteins corresponding to the top 100 hits (E-value < 1 × 10-10) to each query sequence were extracted from the local database using esl-sfetch [89]. Sequences were aligned with MAFFT v7.310 using the E-INS-i strategy and the BLOSUM30 amino acid scoring matrix [90] and trimmed with trimAL v1.4.rev15 using its gappyout strategy [91]. The topologies were inferred using maximum likelihood as implemented in RA×ML v8.2.9 [86] using empirically determined substitution models and rapid bootstrapping (1000 replications). The phylogenies were midpoint rooted and branches with less than 80% bootstrap support were collapsed using the ape and phangorn R packages [92,93]. Phylogenies were visualized using ITOL version 3.0 [88].

To understand the evolutionary histories of SM gene clusters 4 and 10, full-length nucleotide sequences of all 62 strains with short read sequence data were extracted for the entire cluster region (SM gene cluster 4) or the downstream flanking region (SM gene cluster 10) using the previously described SNV analysis procedure followed by Genome Analysis Toolkit’s “ExtractAlternativeReferenceFasta” tool [74]. The resulting nucleotide sequences were aligned using MAFFT v7.310 [90]. Phylogenies were constructed using maximum likelihood as implemented in RA×ML v 8.0.25, using the GTRGAMMA substitution model and rapid bootstrapping (1000 replications) [86]. Phylogenies were midpoint rooted and branches with less than 80% bootstrap support were collapsed. Phylogenies were visualized using ITOL version 3.0 [88].

### Differential metabolite analysis

For natural product analysis, 5 × 10^6^ spores/mL for the indicated strains were grown in 50 mL liquid GMM [94] for five days at 25 °C and 250 rpm in duplicates. Supernatants were extracted with equal volumes of ethyl acetate, dried down and resuspended in 20% acetonitrile (ACN). Each sample was analyzed by ultra-high performance liquid chromatography (UHPLC) coupled with mass spectrometry (MS). The samples were separated on a ZORBAX Eclipse XDB-C18 column (Agilent, 2.1 × 150 mm with a 1.8 μM particle size using a binary gradient of 0.5 % (v/v) formic acid (FA) as solvent A and 0.5 % (v/v) FA in ACN as solvent B that was delivered by a VanquishTM UHPLC system (Thermo Scientific) with a flow rate of 0.2 mL/min. The binary gradient started with 20% B that was increased with a linear gradient to 100% B in 15 min followed by an isocratic step at 100% B for 5 min. Before every run, the system was equilibrated for 5 min at 20% B. The UHPLC system was coupled to a Q Exactive hybrid quadrupole OritrapTM MS (Thermo Scientific). For electrospray ionization, the ion voltage was set at +/−3.5 kV in positive and negative mode, respectively. Nitrogen was used as sheath gas at a flow rate of 45 and as sweep gas at a flow rate of 2. Data analysis was performed using XCMS [54] and Maven [95] software.

## Acknowledgements

A.L.L. was supported by the U.S. National Library of Medicine training grant 2T15LM007450. This work was supported in part by the National Science Foundation (IOS-1401682 to J.H.W. and DEB-1442113 to A.R.), the National Institutes of Health (NIH grant R01 AI065728-01 to N.P.K), and by the Northern Portugal Regional Operational Programme (NORTE 2020) under the Portugal 2020 Partnership Agreement through the European Regional Development Fund (FEDER) (NORTE-01-0145-FEDER-000013 to F.R.). G.H.G. thanks the Fundação de Amparo à Pesquisa do Estado de São Paulo (FAPESP) and Conselho Nacional de Desenvolvimento Cientifico e Tecnológico (CNPq), both from Brazil, for funding and support. This work was conducted in part using the resources of the Advanced Computing Center for Research and Education at Vanderbilt University (http://www.accre.vanderbilt.edu/).

## Author contributions

Conceptualization, A.R., G.H.G., A.L.L.; Methodology, A.L.L., J.H.W, C.L, P.W., J.M.P.; Investigation, A.L.L.; Visualization, A.L.L., J.H.W, P.W.; Resources, G.H.G., F.R., N.P.K., C.L.; Writing, A.L.L, A.R.

---

Figure S1. Alignments showing deletion of genes in SM gene clusters.

Figure S2. Gene phylogenies of the fusarielin-like SM gene cluster 4.

Figure S3. Fusarielin-like gene clusters in Eurotiomycetes and Sordariomycetes.

Figure S4. Novel SM gene clusters in A. fumigatus strains.

Figure S5. Gene phylogenies of SM gene cluster 34.

Figure S6. Two alleles of the idiomorphic SM gene cluster 10 present in one strain each.

Figure S7. Metabolomics analysis of strains with different alleles of the idiomorphic cluster indicates presence of different metabolites.

Figure S8. Idiomorphic locus in other species.

Figure S9. Gene phylogenies of the mobile SM gene cluster 1.

Figure S10. Marker gene phylogenies of all strains and *Aspergillus fischeri*.

Table S1. Summary of strains, sequence data, and assemblies used.

Table S2. All nonsynonymous variants in SM gene cluster genes.

Table S3. Description, locus tags, and gene annotations of all reference Af293 SM gene clusters.

Table S4. XCMS analysis of extracted supernatants from 08-19-02-30,08-12-12-13, and 08-1902-10.

